# Network Propagation-based Prioritization of Long Tail Genes in 17 Cancer Types

**DOI:** 10.1101/2021.02.05.429983

**Authors:** Hussein Mohsen, Vignesh Gunasekharan, Tao Qing, Montrell Seay, Yulia Surovtseva, Sahand Negahban, Zoltan Szallasi, Lajos Pusztai, Mark B. Gerstein

**Affiliations:** Computational Biology & Bioinformatics Program, Yale University, New Haven, CT 06511, USA; Breast Medical Oncology, Yale School of Medicine, New Haven, CT 06511, USA; Yale Center for Molecular Discovery, Yale University, West Haven, CT 06516, USA; Department of Statistics & Data Science, Yale University, New Haven, CT 06511, USA; Children’s Hospital Informatics Program, Harvard-MIT Division of Health Sciences and Technology, Harvard Medical School, Boston, MA 02115, USA; Department of Molecular Biophysics and Biochemistry, Yale University, New Haven, CT 06511, USA; Department of Computer Science, Yale University, New Haven, CT 06511, USA

## Abstract

**Introduction:** The diversity of genomic alterations in cancer pose challenges to fully understanding the etiologies of the disease. Recent interest in infrequent mutations, in genes that reside in the “long tail” of the mutational distribution, uncovered new genes with significant implication in cancer development. The study of these genes often requires integrative approaches with multiple types of biological data. Network propagation methods have demonstrated high efficacy in uncovering genomic patterns underlying cancer using biological interaction networks. Yet, the majority of these analyses have focused their assessment on detecting known cancer genes or identifying altered subnetworks. In this paper, we introduce a network propagation approach that entirely focuses on long tail genes with potential functional impact on cancer development.

**Results:** We identify sets of often overlooked, rarely to moderately mutated genes whose biological interactions significantly propel their mutation-frequency-based rank upwards during propagation in 17 cancer types. We call these sets “upward mobility genes” (UMGs, 28-83 genes per cancer type) and hypothesize that their significant rank improvement indicates functional importance. We report new cancer-pathway associations based on UMGs that were not previously identified using driver genes alone, validate UMGs’ role in cancer cell survival *in vitro*—alone and compared to other network methods—using extensive genome-wide RNAi and CRISPR data repositories, and further conduct *in vitro* functional screenings resulting the validation of 8 previously unreported genes.

**Conclusion:** Our analysis extends the spectrum of cancer relevant genes and identifies novel potential therapeutic targets.

## 1. Background

Rapid developments in sequencing technologies allowed comprehensive cataloguing of somatic mutations in cancer. Early mutation-frequency-based methods identified highly recurrent mutations in different cancer types, many of which were experimentally validated as functionally important in the transformation process and are commonly referred to as cancer driver mutations. However, the biological hypothesis that recurrent mutations in a few driver genes account fully for malignant transformation turned out to be overly simplistic. Recent studies indicate that some cancers do not harbor any known cancer driver mutations, and all cancers carry a large number of rarely recurrent mutations in unique combinations in hundreds of potentially cancer relevant genes [1-7]. These genes are part of a long tail in mutation frequency distributions and referred to as “long tail” genes.

Many long tail mutations demonstrated functional importance in laboratory experiments, but studying them all and assessing their combined impact is a daunting task for experimentalists. This creates a need for new ways to estimate the functional importance and to prioritize long tail mutations for functional studies. A central theme in finding new associations between genes and diseases relies on the integration of multiple data types derived from gene expression analysis, transcription factor binding, chromatin conformation, or genome sequencing and mechanistic laboratory experiments. Protein-protein interaction (PPI) networks are comprehensive and readily available repositories of biological data that capture interactions between gene products and can be useful to identify novel gene-disease associations or to prioritize genes for functional studies. In this paper, we rely on a framework that iteratively propagates information signals (i.e. mutation scores or other quantitative metrics) between each network node (i.e. gene product) and its neighbors.

Propagation methods have often leveraged information from genomic variation, biological interactions derived from functional experiments, and pathway associations derived from the biomedical literature. Studies consistently demonstrate the effectiveness of this type of methods in uncovering new gene-disease and gene-drug associations using different network and score types. Nitsch *et al.* [8] is one of the early examples that used differential expression-based scores to suggest genes implicated in disease phenotypes of transgenic mice. A study by Lee *et al.* shortly followed to suggest candidate genes using similar propagation algorithms in Crohn’s disease and type 2 diabetes [9]. Other early works that use propagation account for network properties such as degree distributions [10] and topological similarity between genes [11-13] to predict protein function or to suggest new candidate genes.

Cancer has been the focus of numerous network propagation studies. We divide these studies into two broad categories: (A) methods that initially introduced network propagation into the study of cancer, often requiring several data types, and (B) recent methods that utilize genomic variation, often focusing on patient stratification and gene module detection (for a complete list, see [14]).

Köhler *et al.* [15] used random walks and diffusion kernels to highlight the efficacy of propagation in suggesting gene-disease associations in multiple disease families including cancer. The authors made comprehensive suggestions and had to choose a relatively low threshold (0.4) for edge quality filtering to retain a large number of edges given the limitations in PPI data availability in 2008. Shortly afterwards, Vanunu *et al.* [16] introduced PRINCE, a propagation approach that leverages disease similarity information, known disease-gene associations, and PPI networks to infer relationships between complex traits (including prostate cancer) and genes. Propagation-based studies in cancer rapidly cascaded to connect gene sequence variations to gene expression changes using multiple diffusions [17], to generate features used to train machine learning models that predict gene-disease associations in breast cancer, glioblastoma multiforme, and other cancer types [18, 19], or to suggest drug targets in acute myeloid leukemia by estimating gene knockout effects *in silico* [20].

Hofree *et al.* introduced network-based stratification (NBS) [21], an approach that runs propagation over a PPI network to smoothen somatic mutation signals in a cohort of patients before clustering samples into subtypes using non-negative matrix factorization. Hierarchical HotNet [22] is another approach that detects significantly altered subnetworks in PPI networks. It utilizes propagation and scores derived from somatic mutation profiles as its first step to build a similarity matrix between network nodes, constructs a threshold-based hierarchy of strongly connected components, then selects the most significant hierarchy cutoff according to which mutated subnetworks are returned. Hierarchical HotNet makes better gene selections than its counterparts with respect to simultaneously considering known and candidate cancer genes, and it builds on two earlier versions of HotNet (HotNet [23] and HotNet2 [24]).

These studies have addressed varying biological questions towards a better understanding of cancer, and they have faced limitations with respect to (i) relying on multiple data types that might not be readily available [17, 18], (ii) limited scope of biological analysis that often focused on a single cancer type [17, 20], (iii) suggesting too many [20] or too few [19] candidate genes, or (iv) being focused on finding connected subnetworks, which despite its demonstrated strength as an approach to study cancer at a systems level might miss lone players or understudied genes [17, 22-24]. To address these issues and parallel the emerging focus on long tail genes and non-driver mutations [2, 4, 5, 25-29], we build on the well-established rigor of propagation and introduce a new approach that particularly prioritizes rarely to moderately mutated genes implicated in cancer. Our analysis spans 17 cancer types and relies centrally on two data types: mutation frequency and PPI connectivity data. We hypothesize that a subset of long tail genes, originally with low mutation frequency ranks, can leverage their positionality in PPI networks and the mutational burden within their extended neighborhoods to play an important role in cancer as signaled by the much higher individual ranks they attain after propagation. These genes can be pinpointed based on their pre- and post-propagation rank differences beyond any subnetwork constraint, and they are described throughout this paper as upward mobility genes (UMGs). To the limits of our knowledge, this is the first propagation approach that focuses entirely on long tail genes.

We efficiently identify a considerable number of UMGs (n = 28-83 per cancer type) and demonstrate their functional importance in cancer on multiple levels. Using somatic mutation data from the TCGA and two comprehensive PPI networks with significant topological differences, STRING v11 [30] and HumanNet v2 [31], we detect UMGs in BRCA, CESC, CHOL, COAD, ESCA, HNSC, KICH, KIRC, KIRP, LIHC, LUAD, LUSC, PRAD, READ, STAD, THCA, and UCEC. These genes reveal a significant number of regulatory pathway associations that would be overlooked when relying on known driver genes alone. Further, *in silico* analysis demonstrates that UMGs exert highly significant effect on cancer cell survival *in vitro* with cancer type specificity, and they outperform genes suggested by other network methods with respect to this impact on cancer cell survival. We then validate a previously unreported subset of the identified genes *in vitro* through siRNA knockout experiments. Finally, we perform a deeper analysis of UMGs’ positionality in a combined STRING-HumanNet v2 PPI network to classify each UMG as a potential cancer driver, drug target, or both. Together with known drivers, we hope that UMGs will draw a more complete portrait of the disease. A python implementation of the approach is available for execution at the cohort or single sample level.

## 2. Results

### 2.1 Overview

First, we generate PPI networks specific to each of 17 cancer types in the TCGA using only genes that are expressed in a given cancer type (Figure 1a). We use the STRING and HumanNet v2 networks that have different topologies and information channels for constructing the networks and use only high quality edges. We then perform propagation over each network, where each sample’s somatic mutation profile includes a quantized positive value ∈ [1,4] for genes with mutations, and 0 otherwise (Figure 1b). Next, we perform hypergeometric test to assess the significance of propagation-based rankings by measuring the enrichment of COSMIC genes above the 90^th^ percentile of ranked post-propagation lists. Results demonstrate high statistical significance across all studied cohorts (*p* < 10^-5^) in a validation of propagation as a tool to rank genes for potential functional importance. We then calculate the difference in pre- (i.e. raw mutation frequency) and post-propagation ranking for each gene. Genes that move up in the rank order in the post propagation list are called UMGs. We construct a preliminary UMG list for each cancer cohort based on stringent final rank cutoff and upward rank increase (i.e. upward mobility) threshold. In this paper, genes whose rank significantly improves during propagation and land in a pre-defined top block of post-propagation ranked lists are retained (Figure 1c). Using this strategy, our approach focuses on long tail genes and excludes frequently mutated genes (including classical cancer drivers) that occupy high ranks before propagation and therefore cannot meet the upward mobility threshold. We identify UMGs separately for each of the 17 cancer types. To further filter UMGs for potential functional importance, we remove genes with minimal or no impact on corresponding cancer cell survival after gene knockdown in the Cancer Dependency Map Project (DepMap) [32]. This step eliminates 4-13% of UMGs (Figure 1d). We finally analyze the biological and topological properties of the shortlisted UMGs on pan-cancer and cancer type levels (Figure 1e).

**Figure 1.**
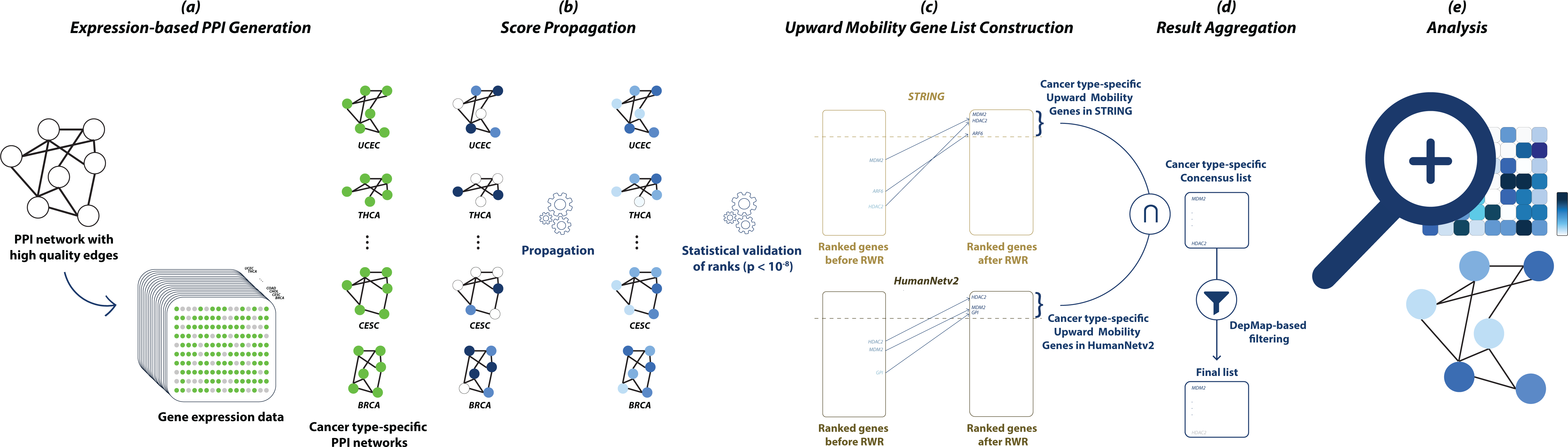
Schematic overview of the UMG identification strategy.

### 2.2 UMGs across 17 cancer types

We report 230 UMGs across 17 cancer types. UMG lists capture the expected biological heterogeneity of cancer types: 76 genes (33%) are specific to one cancer type, 116 (50.4%) to 2-9 types, and only 38 (16.5%) to 10 or more types. The longest list of UMGs corresponds to CESC (n = 83 genes) and the shortest to CHOL (n = 28). Hierarchical complete linkage clustering of cancer types (right of Figure 2) using UMG list membership and DepMap dependency scores of the genes (which reflect their importance in cell growth) reveals interesting patterns. Similar to results based on driver gene sets identified in [7], subsets of squamous (ESCA, HNSC, and LUSC) and gynecological (BRCA, CESC, and UCEC) cancers cluster together. Close clustering results also correspond to the lung (LUAD and LUSC) and colon and rectum (COAD and READ) as tissues of origin, while others match with the rates of driver mutations across cancer types (i.e. Figure 1D in [7]), particularly (i) STAD and CESC, (ii) KIRP, READ, and COAD, and (iii) LUSC, LUAD, HNSC, ESCA, and LIHC, suggesting similarities between driver and long tail mutational patterns. Interestingly, UMGs specific to a single cancer type (left of Figure 2) include a considerable number of genes whose products have similar functions such as *COL4A1* and *COL1A1* that encode different types of collagen (specific to ESCA), and triplets of genes that encode proteins in the 26S proteasome complex (*PSMC1/2/3*, specific to UCEC) and mitogen-activated kinases (*MAPK1* and *MAP2K1/2*, specific to THCA). Functional gene clusters shared among cancer types include *DYNC1LI2/I2/H1* that encode different components of the cytoplasmic dynein 1 complex and *PPP1CC/1CA/2CB/2CA* that encode subunits of protein phosphatase enzymes. The circos plot [33] of Figure 2 shows the distribution of UMGs across cancer types, their relative ranks within UMG lists, and their impact on cancer type-specific cell survival.

**Figure 2.**
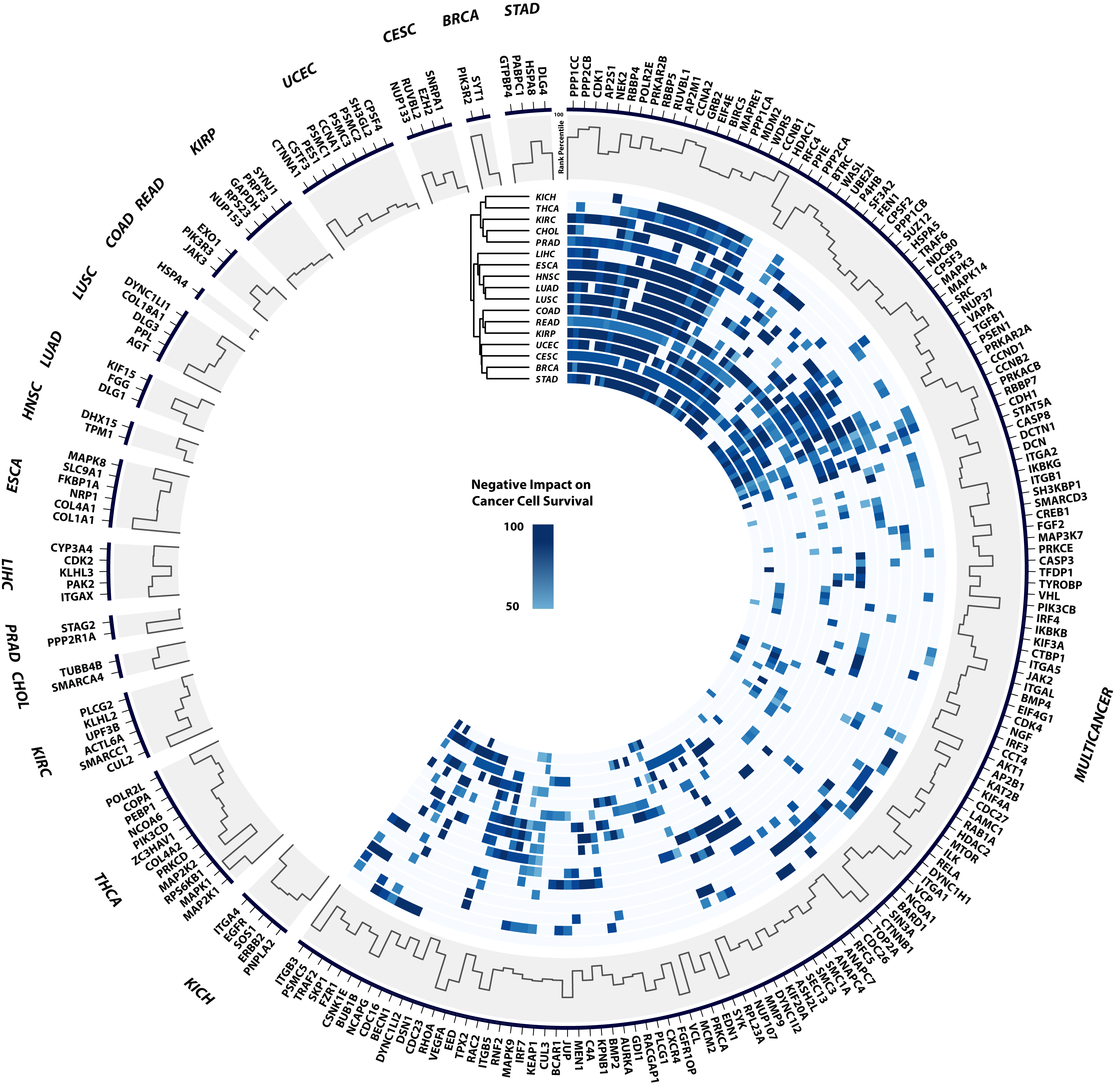
Distribution of UMGs across 17 cancer types. Right: genes in 2 or more cancer types. Dendrogram is based on hierarchical clustering of heatmap rows. Each heatmap value corresponds to a percentage-based score of a cancer type’s cell lines whose survival is negatively impacted by a gene’s knockout. For each value, the maximum percentage across RNAi and CRISPR experiments is selected. Left: cancer type-specific genes. Histogram throughout the plot corresponds to the normalized rank of each UMG in the lists it belongs to.

### 2.3 UMGs reveal known and novel cancer-pathway associations

Biological enrichment analysis of UMGs, separately and in combination with known drivers, confirms already known functional importance of the UMGs and suggests new associations between cancer types and biological pathway alterations. UMGs have statistically significant associations (Benjamini *p-adjusted* < 0.05) with most of the oncogenic pathways curated by Sanchez-Vega *et al.* (8 of 10) [34], alone and also when combined with cancer type-specific drivers (Figure 3a). These results indicate that UMGs are members of known biological pathways and can broaden the study of biological processes that contribute to malignant transformation. This is particularly relevant in cancers where driver gene-based pathway associations revealed only a few relevant pathways (e.g. KICH and CHOL in [7]). Interestingly, the p53 pathway has only a small number of associations with UMGs in contrast to more associations we detected with the cell cycle, TGF-beta and Hippo signaling pathways. Other known cancer pathways are also altered by UMGs and include Notch, HIF-1 and mTOR. Notably, the number of cancer type-specific pathway associations does not correlate with the size of UMG lists. For example, KICH, which has one of the smallest lists of UMGs (n = 41 genes), has a sizeable set of pathway associations, while CESC with the largest UMG list (n = 83) has considerably fewer associations. These findings suggest greater diversity in altered biological processes that lead to development of KICH compared to CESC.

**Figure 3.**
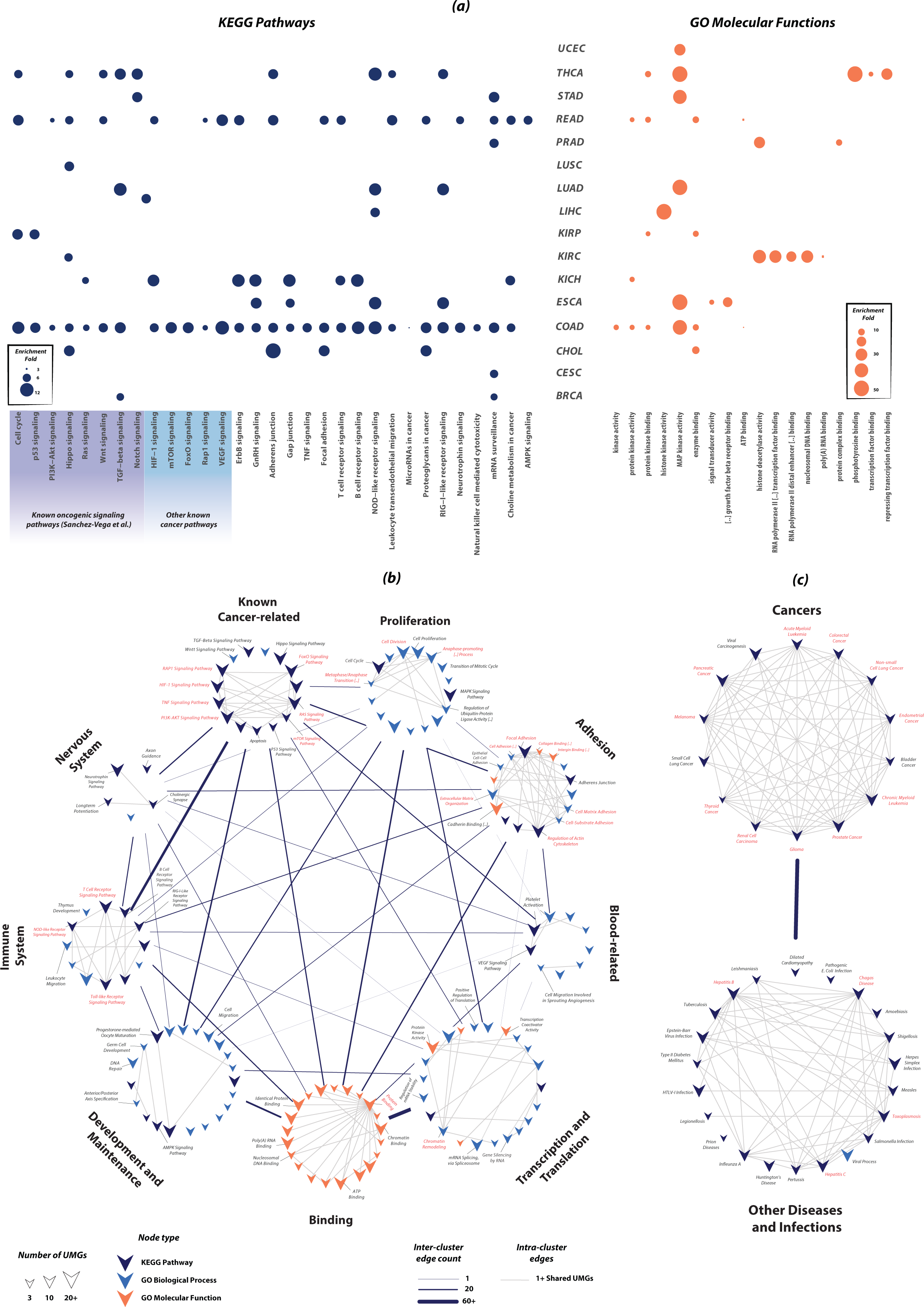
Biological enrichment results for UMGs at cancer type and pancancer levels. (a) UMGs uncover known and novel associations between cancer types and biological pathways. Enrichment analyses are performed for each cancer type’s combined list of UMGs and drivers. Shown results correspond to significant pathway and molecular function associations exclusively uncovered by UMGs. (b) Pancancer analysis of all 237 UMGs visualized using EnrichmentMap allows for the identification of biological pathways, processes and functions strongly associated with UMGs (in red) that suggests potential therapeutic targets. (c) Similar analysis to (b) on clusters of KEGG mega-pathways uncover disease-disease and disease-infection associations driven by UMGs

On the pancancer level, we partitioned enrichment results for all 230 UMGs into 9 clusters based on biological function (Figure 3b). Using EnrichmentMap (EM) [35], we built a network of intra- and inter-cluster similarity measured through gene overlap between enrichment entities (i.e. pathways, biological processes and molecular functions; Methods). Connectivity patterns within the EM network provide insights into the sets of entities and UMGs. Within 6 of the 9 clusters, namely ones with known relation to cancer pathways, proliferation, adhesion, binding, immune response and transcription and translation, we identified biological entities with high connectivity (red labels, Figure 3b). These entities include oncogenic pathways such as PI3K-AKT, RAS, and mTOR, and important biological processes including cell matrix adhesion and chromatin remodeling. Underlying their high connectivity is a selected subset of UMGs with high frequency in their constituent edges (Table 1). Susbsets of these frequent UMGs encode subunits of proteins (and protein complexes) with strong association with cancer such as *MDM2*, *PIK3R2/R3/CB/CD*’s products in phosphatidylinositol kinases (PI3Ks) [36], and *IKBKB/G* with regulatory subunits in an inhibitor of the Nuclear Factor Kappa B kinase (NFKB) [37]. Given their significant and wide range of biological functionality, these genes constitute a potential subset of potent drug targets. A similar analysis on KEGG mega-pathways corresponding to diseases and infections revealed another subset of frequent UMGs and demonstrated the ability of long tail gene analysis to uncover disease-disease/infection associations (Figure 3c, Table 1). Observed associations include well-studied ones between multiple cancers and Hepatitis C [38], Type II Diabetes Mellitus [39, 40], and HTLV-I infection [41].

**Table 1.**
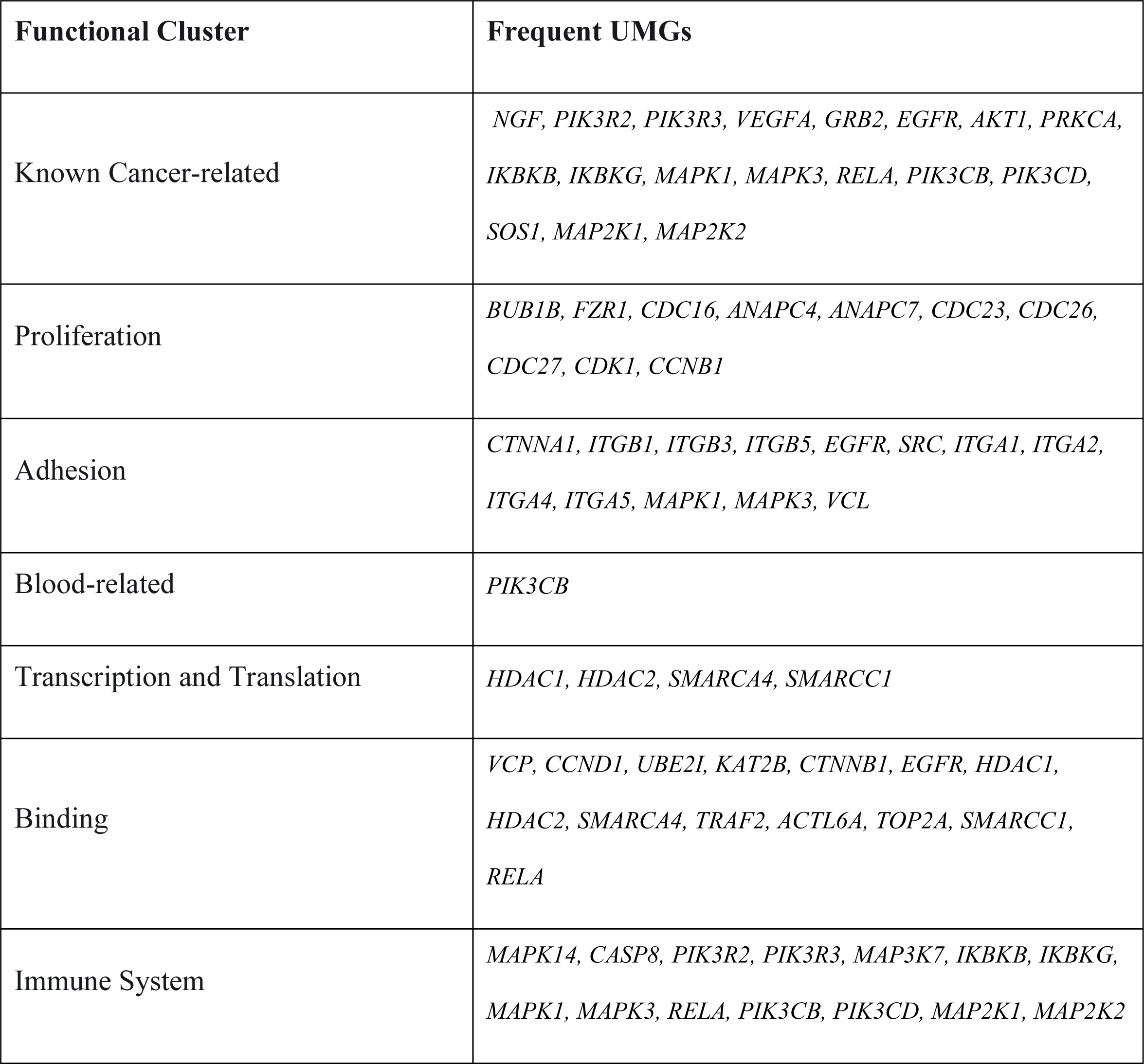
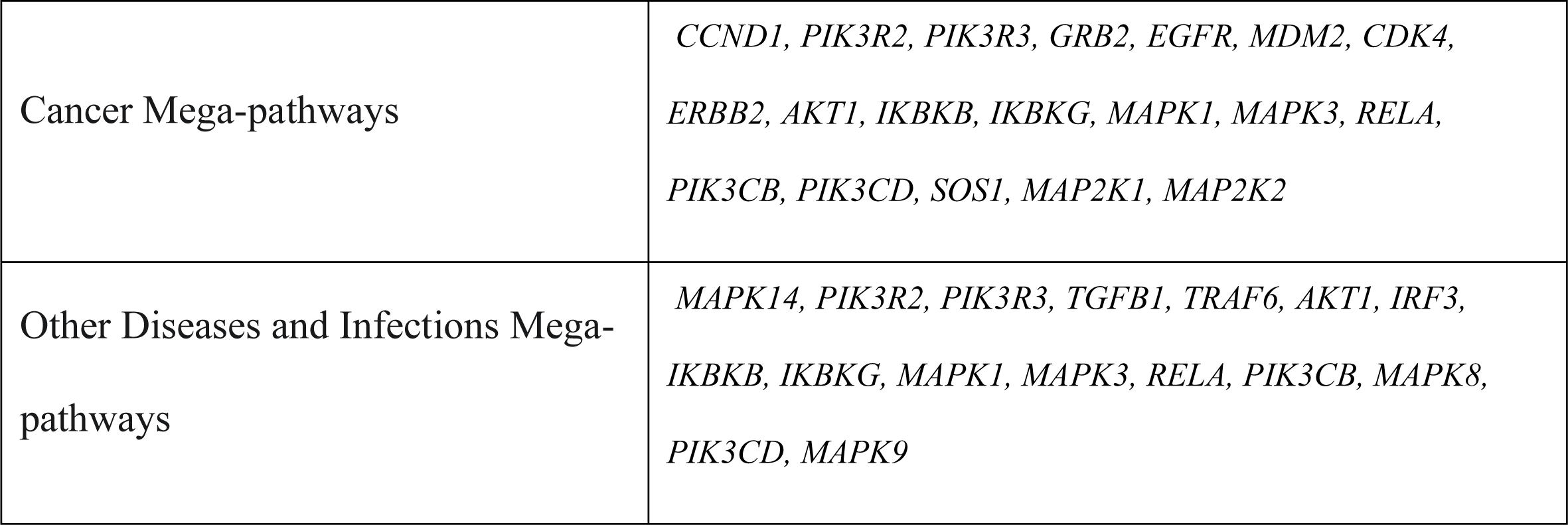
Frequent UMGs within EnrichmentMap functional clusters

### 2.4 UMGs impact survival of cancer cells in vitro

To assess the functional importance of UMGs in cancer cell survival *in vitro*, we obtained their cancer type-specific dependency scores from the DepMap project. DepMap reports results on comprehensive genome-wide loss of function screening for all known human genes using RNA interference (RNAi) and CRISPR to estimate tumor cell viability after gene silencing in hundreds of cancer cell lines [32]. A dependency score of 0 corresponds to no effect on cell viability, and a negative score corresponds to impaired cell viability after knocking down the gene; the more negative the dependency score, the more important the gene is for cell viability. We used the most recent data release that accounts for batch and off-target effects and therefore provides more accurate estimates of functional impact [42].

We found that cancer type-specific mean scores of UMGs’ negative impact on the survival of cancer cell lines is higher (i.e. more negative) than non-UMGs’ across all 17 cancer types, and in both CRISPR and RNAi experiments. The knockout of UMGs consistently yields a stronger negative *in vitro* effect on the survival of more cell lines than that of non-UMGs (Mann-Whitney U test, *p* < 5 ×10^-3^, Methods).

UMG detection is entirely focused on prioritizing long tail genes to parallel the recently growing research on the topic. Most existing network methods have understandably been designed to focus on uncovering known cancer genes or geared towards other goals—such as detecting subnetworks that maximize coverage of mutational profiles or are highly mutated. To have a better understanding of the specifications of UMGs, we still compared their impact on the survival of cancer cell lines to that of genes selected by two state-of-the-art propagation methods, FDRNet [43] and Hierarchical HotNet (HHotNet) [22]—in 3 different settings, and nCOP [44], a non-propagation method that recently demonstrated an ability to uncover non-driver genes across multiple cancer types. HHotNet reported statistically significant results after the integration over both PPIs in only 5 out of the 17 cancer types. Hence, we included two other settings (largest and all subnetworks) where the method was able to report statistically significant results in one of the PPIs. FDRNet results successfully generated results on the STRING PPI, and hence its reported results across cancer types are based on this PPI (Methods).

In terms of DepMap scores, almost all methods’ selected gene sets have negative impact on cell survival. Of the methods we tested for gene selection, the UMGs have the strongest negative impact on cancer cell survival across all cancer types in both CRISPR and RNAi experiments (Figure 4a). Similarly, the median percentage-based score of cell lines negatively impacted by UMGs’ knockout is consistently higher than that for genes selected by other methods in 28 out of the 32 cancer type-assay combinations (Figure 4b), with the remaining 4 including 3 ties. Notably, a number of UMGs have an extremely strong negative impact on cell survival across cancer types. For instance, PRAD, READ, and THCA sets include genes with mean DepMap CRISPR score < -2 in their cell lines, and all other cancer types except HNSC include genes with score < -1.7. Similar results were also obtained for these comparisons before the optional DepMap filtering step that only removed 4-13% of UMGs. As FDRNet, HHotNet and nCOP do not solely focus on long tail genes, we performed the same experiments after removing known cancer-specific driver genes from all gene lists (including UMGs’) during comparisons. Similar results were also obtained.

**Figure 4.**
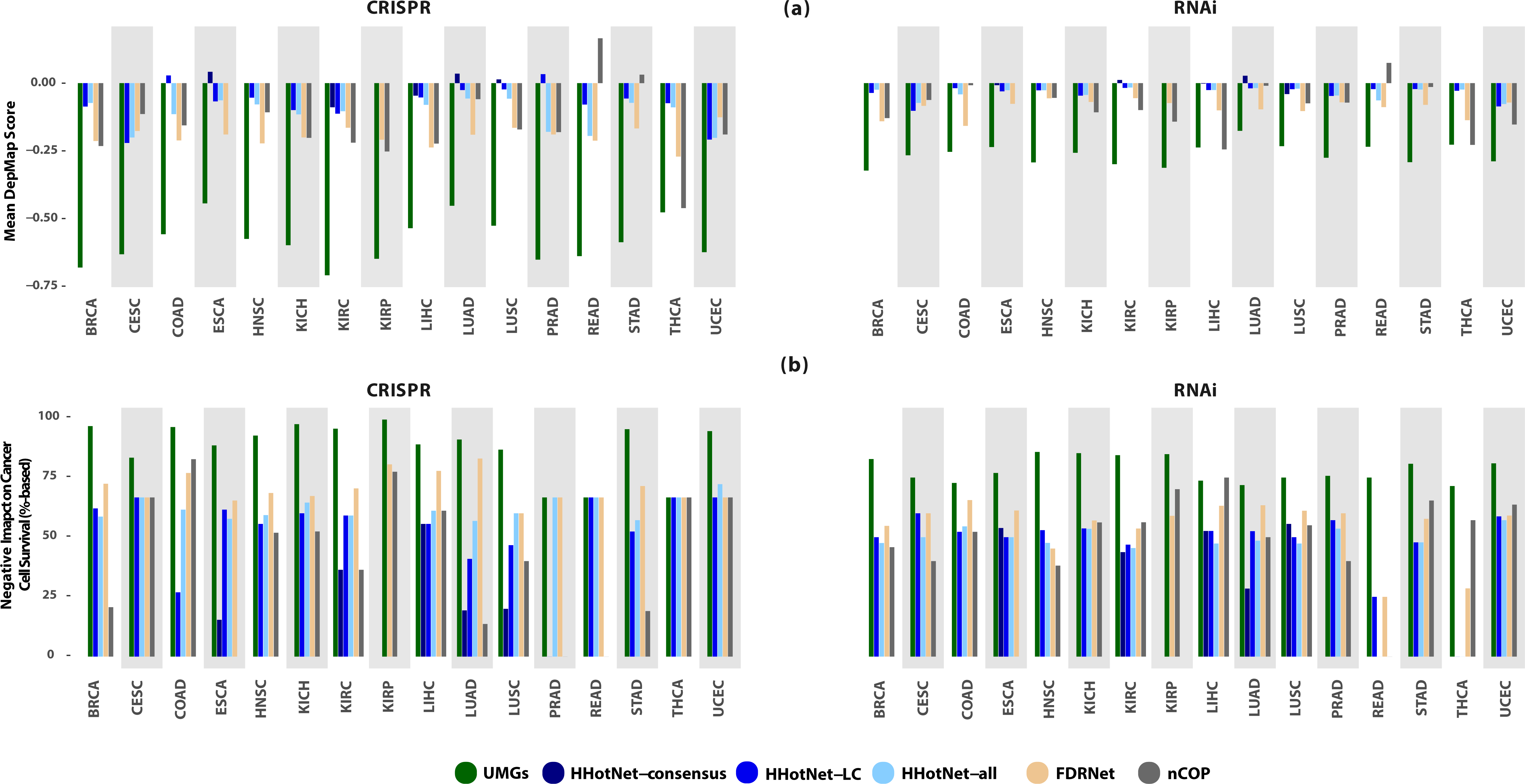
Comparisons with other methods. (a) UMGs demonstrate considerably stronger (CRISPR- and RNAi-measured) impact on survival of cancer cell lines than other non-driver genes suggested by HHotNet (in 3 settings) and nCOP. Higher negative values indicate greater negative effect on cell survival after gene knockdown. (b) UMGs’ strong impact on the survival of cancer cell lines is significantly broader than that of genes selected by HHotNet and nCOP. The median percentage-based score of cancer cell lines negatively impacted by UMGs’ knockout is consistently higher with cancer type specificity.

### 2.5 UMGs as “weak drivers” and potential novel drug targets

The aim behind identifying UMGs is to expand the narrative of known driver genes underlying cancer in line with many recent studies whose results defy the neutrality of long tail genes and passenger mutations in carcinogenesis [2, 4, 5, 25-29]. In this section, we categorize each UMG as a potential “weak driver” that supplements known drivers during carcinogenesis, a potential drug target whose suppression kills cancer cells, or both, according to its positionality in PPI networks with respect to currently known drivers.

In the propagation framework we use, two of the most important factors that determine a node’s score after convergence are the number of high scoring nodes within its neighborhood and the connectivity of these neighbors. For a node to rank higher, the best case scenario involves having near exclusive connections with multiple neighbors (*k* ≥ 1 steps) whose initial score is high. We study these properties of each cancer type’s UMGs. We use a composite PPI network that merges signals from STRING and HumanNet v2 by including the union of high quality edges of both networks. Figure 5 shows a representative network that corresponds to BRCA, with all others included in the supplement. For convenience in visualization, we include immediate neighborhoods of each node and UMG-driver edges only.

**Figure 5.**
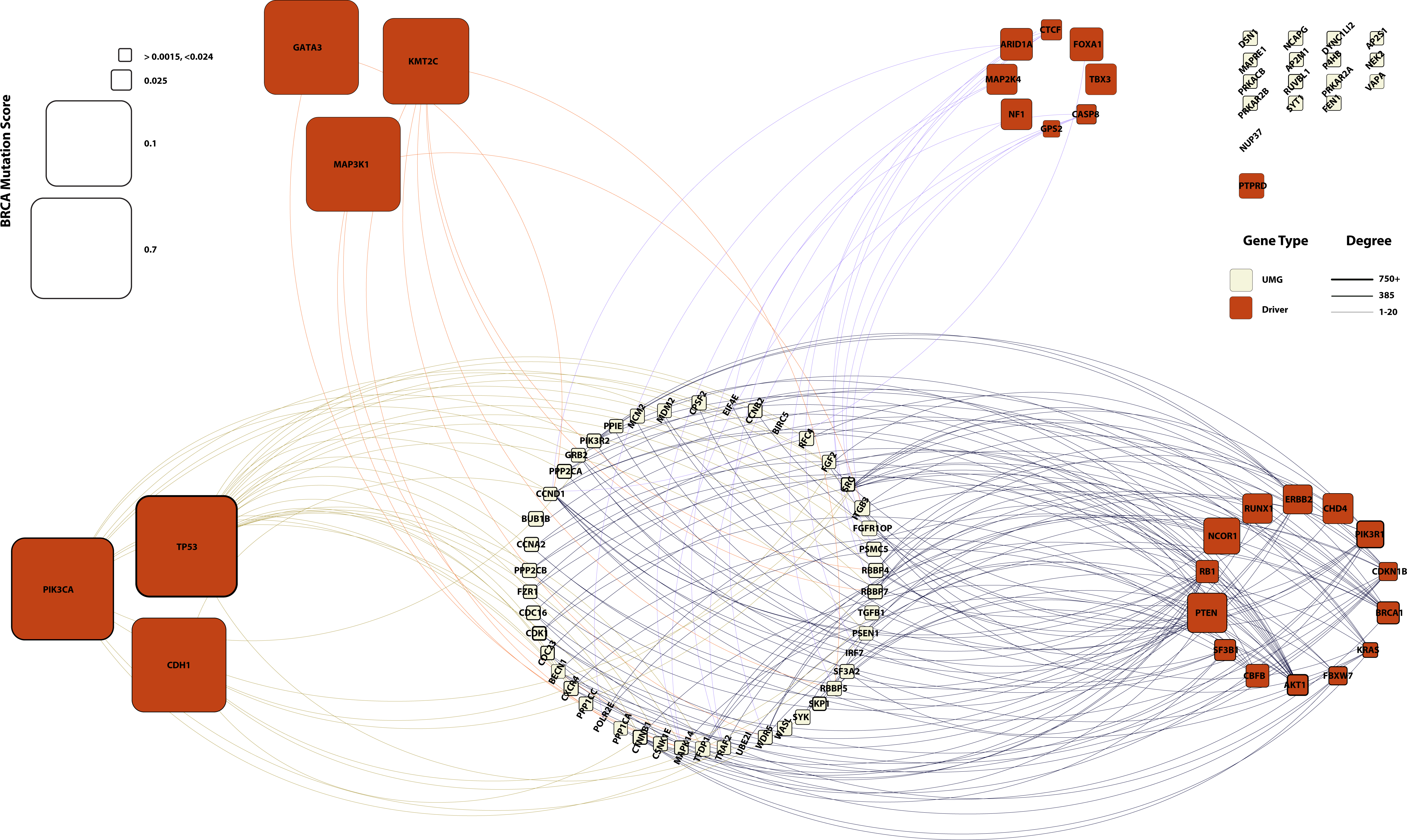
PPI network analysis of the relationships between UMGs (white nodes) and known driver genes (red) in breast invasive carcinoma (BRCA) suggest roles of UMGs. Driver genes are split into categories based on initial mutation score and node degree: (i) high score, high degree (bottom left), (ii) high score, low degree (top left), (iii) low score, low degree (top right) and (iv) low score, high degree (bottom right). UMGs connected to driver subsets (i) and (ii) (olive and orange edges) and ones with no mutation score (e.g. *POLR2E*) are likely to be drug targets. UMGs connected to (iii) and (iv) and ones without connections to drivers (top right corner, e.g. *DSN1*) are likely to be “weak drivers.”

The first category of UMGs includes ones connected to high scoring drivers (Figure 5 left side, olive and orange edges). By virtue of sharing connections with these potent and frequently mutated drivers, this subset of UMGs likely includes cancer type-specific potential drug targets with little effect on carcinogenesis. This becomes even more relevant for UMGs connected to high degree, high scoring drivers (via olive edges). Building on the same reasoning, low scoring drivers might not be the dominating force driving cancer across the majority of samples. UMGs connected to these low scoring drivers (Figure 5 right side, dark blue and purple edges) constitute the second category and are likely to have a supplementary driving force. Interestingly, the third category includes an often small subset with nearly no observed mutations in the cohort (e.g. 6 genes in Figure 5: *NUP37*, *UBE21*, *POLR2E*, *IRF7*, *BIRC5*, and *EIF4E*). Such genes are likely to be drug targets or false positives limited by the size of the cohort under study. The fourth category includes UMGs with positive initial score and no connections to driver genes (Figure 5, top right grid). These genes’ positive scores and connectivity with non-drivers significantly lift their rank during propagation and render them potentially overlooked weak drivers. While most UMGs are designated either potential drug targets or weak drivers, others are connected to multiple types of driver genes and accordingly might be considered for both (e.g. *RBBP5* with multi-colored edges in Figure 5).

### 2.6 UMGs bridge gaps in literature and suggest novel genes

The study of cancer has long been interdisciplinary, often in the realms of various scientific and medical spheres. Disciplinary paradigms evolved over time to produce varying types of associations between genes and cancers. To further support the functional importance of UMGs, we manually cross referenced our UMG lists with publications and found that a large percentage of UMGs have been reported to play a role in cancer based on functional experiments. This percentage is as high as 85% of UMGs in cancer types like BRCA. Surprisingly, the same percentage drops to only 31% when we used CancerMine v24 to find literature-based associations [45]. CancerMine is an automated tool that applies text mining on existing literature to report drivers, oncogenes, or tumor suppressors across cancers. Similar results were obtained across cancer types.

### 2.7 Screening experiments validate 8 new genes in vitro

We then conduct a series of siRNA knockdown experiments to further investigate the functional importance of the UMGs without any previously reported functional validation with respect to their effect on cancer phenotype. The majority (n = 28) of those UMGs are selected alongside a number of functionally related genes (Methods). Experimental results further highlighted the significance of 8 genes, 4 UMGs and 4 closely related ones on the functional level. The knockdown of these genes significantly impacted the survival of (1-3 out of 4) cancer cell lines based on the threshold of 3 standard deviations with respect to negative control samples. Validated UMGs are namely *POLR2L, POLR2E, DCTN1,* and *PSMC3*, with the remaining 4 genes being (i) *RPS10* which encodes a ribosomal protein, and (ii) *POLR2I*, *POLR2G*, and *POLR2H* which encode subunits of RNA polymerases—similar to validated UMGs *POLR2L* and *POLR2G*. Results of *in vitro* experiments significantly expand the potential of UMGs to suggest novel cancer-associated genes and drug targets.

## 3. Discussion

In this paper, we expand the set of propagation approaches to parallel the growing interest in long tail genes in cancer. We introduce a computationally efficient approach based on the notion of upward mobility genes that attain significant improvements in mutation score-based ranking after propagating through PPI networks. By virtue of high post-propagation ranks, cancer-related biological function, and significantly strong impact on cancer cell line survival, our approach prioritizes long tail genes across 17 cancer types. To reduce false positivity rate, we integrate results over two major PPI networks, filter out nodes whose genes are unexpressed in each cancer type’s tumor samples, and statistically validate rankings and cell survival impact.

Biological analysis of UMGs provides novel insights and demonstrates strong correlations with studies performed on known cancer drivers. Enrichment analysis results unlock a wide range of potential associations between key pathways and cancer types. A network-based analysis of enrichment results allows for classifying UMGs based on their centrality to biological functions, opening the door for a more informed drug targetability. Another network-based analysis categorizes each UMG as a “weak driver,” cancer type-specific drug target. Manual curation of literature further validates UMGs’ connection to cancer which could be overlooked by automated literature mining alone.

Results suggest that we have not reached a point of data saturation with respect to analyzing long tail genes yet. The generation of new datasets will likely improve results in rare cancer types such as cholangiocarcinoma (CHOL) and chromophobe renal cell carcinoma (KICH). Novel discoveries on frequently mutated genes in cancer, among which are many drivers, will likely reflect on the study of long tail genes as well. This was particularly evident in our PPI positionality analysis: with 3 or less drivers identified in KICH and READ by Bailey *et al.* [7], most of these cancer types’ UMGs belong to the third and fourth categories (near-zero mutation scores and no connections with drivers). Another example is CHOL, with its small cohort that brings most UMGs into the third category (no observed mutations). Finally, we note that bridging gaps across disciplines is essential to biomedical knowledge production. The oncogenic validation of potential drug targets in UMGs also remains central to changing their status from potential to clinically actionable ones.

## 4. Methods

### 4.1 Availability

UMG detection code is available at https://github.com/gersteinlab/UMG.

### 4.2 Somatic mutation data

The results in this paper are in whole or part based upon data generated by the TCGA Research Network: https://www.cancer.gov/tcga. Variants from the MC3 high quality somatic mutation dataset (n = 3.6 M) [46] are used to generate initial scores for each of the 17 cancer types. Sample-gene matrix for each cancer type includes mutation counts restricted to splicing and coding exonic variants based on RefSeq hg19 annotations by ANNOVAR 2018b [47]. Each count is normalized by gene length values provided by bioMart Bioconductor package [48]. Each non-zero value is then converted to a discrete number in [4 4] based on its position with respect to 50^th^, 70^th^ and 90^th^ percentiles in the cancer type-specific normalized mutation frequency distribution. Gene ranks before and after propagation are calculated based on the mean frequency within each cohort.

### 4.3 PPI networks

STRING v11 and HumanNet v2 functional network (FN) are downloaded from https://string-db.org/ and https://www.inetbio.org/humannet/, respectively. We perform edge filtering on both PPIs and retain edges with a confidence score equal to or higher than 0.7 in STRING and the top 10% of edges in HumanNet v2. The networks after this filtering have |*V*| = 17,130 and 11,360 vertices and |*E*| = 419,772 and 37,150 undirected edges, respectively. We then generate cancer type-specific PPI networks by selecting the largest connected component in each network and filtering out genes unexpressed in the tumor samples of each cancer type.

### 4.4 Expression-based filtering

Gene expression filtering is performed on TCGA expression data corrected for study-specific biases and batch effects from RNASeqDB [49]. For each cancer type, genes with FPKM > 15 in > 20% of tumor samples are retained in the cancer type specific PPI network.

### 4.5 Propagation score calculation

To calculate propagation scores, we use an approach that imitates random walk with restart. Briefly, let the PPI network be represented as *G* = (*V*, *E*), where *V* is the set of gene products and *E* is the set of edges. Further, let *W* be the weighted adjacency matrix of *G*. We choose to normalize *W* such that *W’* = *W* . *D^-1^*, where *D* is the diagonal matrix of columns sums in 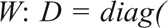 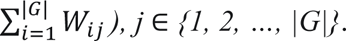

Let *M* be a |*G*| × *N* matrix with somatic mutation profiles of *N ≥* 1 samples over genes from which *G*’s nodes originate before transcription. *Sij* is a positive value for each *gi* ∈ *G* with mutations in sample s*j* ∈ *S*, and 0 otherwise. Propagation is then executed within each sample until convergence according to the following function:

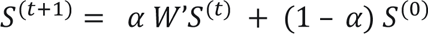

where *S^(0)^ = M* and *α* ∈ [0.5, 1]. Convergence of this propagation technique is guaranteed. We summarize the proof noted in [50] below for the sake of completeness.

The function above can be written at convergence as *S* = *VS + (1 – α) S^(0)^, where V = α W’*, which can also be rearranged into *S* = *(1 – α) (I - V)^-1^ S^(0)^*. For convergence to a unique, non-negative solution to be guaranteed, *(I - V)^-1^* > 0 must hold.

*Lemma 1.* Largest eigenvalue of V < 1. *W’* is a column-stochastic matrix. Per the Perron-Frobenius theorem, its eigenvalues ∈ [-1, +1]. Since *α* < 1, the largest eigenvalue (i.e. spectral radius) of *V* < *α <* 1.

*Lemma 2. (I - V)^-1^* exists, and is non-negative. (*I – V*) is an M-matrix since its in the form *sI* – *B*, with *s* = 1 > 0, *s* >= largest eigenvalue of *B* (i.e. *V*) by Lemma 1, and *V* > 0. An M-matrix is inverse positive, hence (*I - V)^-1^* > 0.

Convergence can also be achieved iteratively [51, 52], which we apply and is more commonly deployed with large PPI matrices for practical considerations. The value of *α* we pick is 0.8. Other values in the [0.6, 0.8] range have little effect on results.

### 4.6 Upward mobility gene identification

The mobility status of a gene is determined by its rank before and after propagation. A gene’s rank is calculated according to its arithmetic average score across samples. For each gene *gi* ∈ *G*,

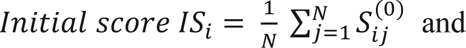

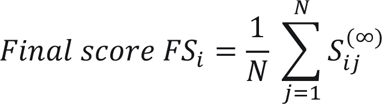

Let *RIS* and *RFS* be the lists of gene ranks in IS and FS, respectively, i.e. *RISi* = rank of *gi* in sorted *IS* and *RFSi* = rank in sorted *FS.* The mobility status of *gi*, *MSi*, is then calculated as the difference between *RISi* and *RFSi* as:

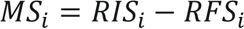

Since higher scores lead to a higher rank, and a higher rank has a lower value (i.e. rank 1, 2, … |*G*|), genes whose ranks improve because of propagation have positive MS values, and ones with lowered ranks (downward mobility) negative ones.

We then define upward mobility status according to two parameters: mobility *β* and rank threshold *T*. 

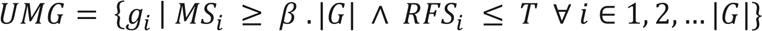

Mobility *β* value determines the minimum upward jump size a gene needs to make to be considered for UMG status. For instance, a *β* value of 0.1 in a PPI network with 10,000 nodes requires a gene’s position to improve by a minimum of 1,000 ranks. We choose stringent values of *β* dictated by TCGA cohort size and the variance of each cancer type’s mutational. Cancer types with a high number of samples and/or a high variance of gene mutation frequency receive a value of 0.25 (BRCA, COAD, HNSC, LUAD, LUSC, PRAD, STAD, UCEC), others with moderate variance a value of 0.2 (CESC, KIRC, KIRP, LIHC) and 0.15 (ESCA, READ), and low variance and/or cohort size cancer types a value of 0.05 (CHOL, KICH, THCA). These values ensure that to be considered a UMG, a gene has to jump hundreds to thousands of ranks during propagation depending on the PPI network and cancer type under study. Rank threshold *T* specifies the minimum rank a gene needs to achieve after propagation to be considered a UMG. We choose *T* = 1,000 to strictly focus on the top 10-16% of genes (i.e. approximately top 10% in STRING and top 16% in HumanNetv2), a threshold that has proved to be effective in other studies [20].

We further apply two optional selection criteria on the final UMG lists based on (i) each gene’s DepMap scores in CRISPR and RNAi experiments and (ii) propagation within multiple PPIs. Per (i), UMG becomes:

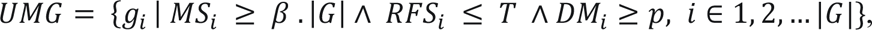

where *p* is the proportion of cancer type-specific cell lines in which a gene’s DepMap score is negative (i.e. its knockout has negative impact on cancer cell survival), and *DMi* is the maximum value across CRISPR and RNAi experiments. We choose *p* = 0.5 (50%), which ends up eliminating 2-10 genes out of 30-91 genes per cancer type. Per (ii), integration of lists across *K* PPI networks yields the intersection of lists. In this paper, to increase confidence is selected genes, we integrate lists over cancer type-specific STRING and HumanNet v2 networks. Formally,

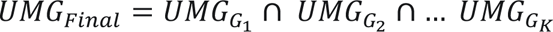

### 4.7 Statistical validation of rankings

To assess the validity of ranking after propagation, we tested if known COSMIC genes [53] are ranked significantly higher than other genes using the hypergeometric statistical test as earlier applied in [20]. Results show strong enrichment of COSMIC genes above the 90^th^ percentile of ranked genes for both PPI networks (*p* < 10^-5^ across all cancer types).

### 4.8 Driver and COSMIC genes

Cancer type-specific driver genes were obtained from Beiley *et al.*’s except for COAD and READ which were combined into a single group in that study. For these two cancer types, we designated tissue-specific COSMIC v90 genes as the driver genes.

### 4.9 UMG vs non-UMG comparisons

In the first set of comparisons, Mann Whitney U one-sided test is used to compare the distribution of a percentage-based score of negatively impacted cell lines by UMGs *vs* non-UMGs in each cancer type. Each gene’s percentage-based score value is equal to the percentage of its negative DepMap scores among *k* cancer type-specific cell lines and the average of these values (to account for distribution of DepMap scores across cell lines). To calculate a more stringent score and reduce false positives, we also assume the presence of at least one cancer cell line with a non-negative DepMap score, which especially accounts for cancer types with a small number of cell lines in the DepMap database. Hence, the score is the sum of each gene’s *k* + 1 values mentioned above divided by *k* + 2. Alternative hypothesis for each of the Mann Whitney U tests is *H*_1_ = *ψ*(*UMG*) is shifted to the right of 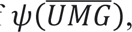, where *ψ*(*X*) is the percentage-based distribution of negatively impacted cell lines over genes in set *X)*. Cancer type-specific cell lines are selected based on annotations provided in the DepMap dataset. For cancer types not represented among the cell lines in DepMap, we used values across all 750 (CRISPR knockout data) and 712 (RNAi) cell lines. A negative DepMap dependency score indicates decreased cell survival after gene knockout in a particular cell line. For RNAi experiments, we use DEMETER2 data with enhanced batch and off-target processing as described in [42].

### 4.10 UMGs vs gene candidates identified by other network methods

Hierarchical HotNet (HHotNet) generates statistically significant results (*p* < 0.05) in only 5 of the 17 cancer types after integrating its results for both PPI networks (HHotNet-consensus): ESCA, KIRC, LIHC, LUAD and LUSC. As a result, we include HHotNet results from two other settings described below. In 13 cancer types, HHotNet generates statistically significant results for one of the two PPI networks, and in two others (PRAD and READ) significant result with a relaxed threshold (0.05 < *p* < 0.1). We include HHotNet results from both the largest subnetwork (HHotNet-LC) and all subnetworks with more than one node (HHotNet-all) in comparisons. Namely, for 15 cancer types, we choose results from STRING in BRCA, ESCA, HNSC, KICH, KIRC, LIHC, LUAD, LUSC, STAD and THCA and from HumanNet v2 in CESC, COAD, PRAD, READ, and UCEC. In CHOL and KIRP, HHotNet results were not statistically significant for both PPI networks, so we exclude results for this method. In all runs, we execute HHotNet in default settings with 1000 permutations using the second controlled randomization approach suggested in [22]. In FDRNet, we run the method to detect subnetworks for all seed genes and in default settings. We obtain MutSigCV2 [54] *p*-values across TCGA cohorts from http://gdac.broadinstitute.org/ and convert them to local FDR values using the scripts provided by FDRNet. We use FDRNet results for 16 cancer types over the STRING network as this method was not able to detect any subnetwork over HumanNetv2 for almost all seed genes (664/673, 98%). No FDRNet results could be produced for CHOL. In nCOP, we use lists of rarely mutated genes reported in [44] (Figure 4) on the TCGA somatic mutational dataset in 15 of the 17 cancer types studied in our paper (all except CHOL and ESCA). As HHotNet and nCOP do not primarily focus on long tail genes, we remove driver genes from these methods’ gene lists to ensure balanced comparisons with UMGs. It is worth noting however that including driver genes or the small percentage of UMGs filtered in the last step of the pipeline did not have a considerable impact on results.

### 4.11 Enrichment analysis

Enrichment analysis to identify KEGG Pathways and GO molecular functions and biological processes is performed on DAVID v6.8 [55]. DAVID chart results with Benjamini *p*-adjusted < 0.05 are selected for analysis. Network visualization is executed using EnrichmentMap v3.0 on Cytoscape v3.8.2 [56], with a comprehensive subset of results related to cancer shown in Figure 3. Frequent terms highlighted in red in Figure 3b have ≥ 5 intra-cluster edges and those in Figure 3c ≥ 10 edges. Frequent UMGs in Table 1 are identified based on their presence edges between highlighted nodes according to the same thresholds (i.e. ≥ 5 and ≥ 10).

### 4.12 PPI analysis

Composite PPI is the union of high quality edges in STRINGv11 and HumanNet v2. Initial score of each gene is the one based on somatic mutations across a cohort as described earlier. Drivers are split according to initial score and degree with thresholds of 150 and 0.075, respectively. Initial scores of < 0.0015 are zero-fied to attain lower FPR. Visualization and degree calculation is executed using Cytoscape v3.8.2.

### 4.13 Experiment validation: siRNA screening and annotation

Cell lines (MDAMB231, MDAMB468, BT549, HCC187) were cultured in RPMI medium supplemented with 10% HI-FBS and penicillin/Streptomycin (1:100). siRNA transfection experiments were performed at the Yale Center for Molecular Discovery. Reverse transfections were performed using 384-well tissue-culture treated plates (Corning CLS3764) pre-plated with siRNAs to achieve 20 nM final assay concentration. RNAiMax transfection reagent (Invitrogen) was added to plates according to manufacturer’s recommendations and incubated with siRNAs for 20 minutes. Cells were then seeded at plating densities optimized during assay development (MDAMB468, HCC1187, and BT549 seeded at 4000 cells per well; MDAMB231 seeded at 1000 cells per well). After 72 hours, CellTiter-Glo (Promega) kit was used to monitor viability. Each screening plate contained 16 replicates of both negative siRNA controls (siGENOME Smart Pool non-targeting control #2, Dharmacon) and positive siRNAs controls (siGENOME Smart Pool Human PLK1 or KIF11, Dharmacon). Signal-to-background (S/B), coefficient of variation (CV) and Z prime factor (Z’) were calculated for each screening plate using mean and standard deviation values of the positive and negative controls to monitor assay performance. For each cell line, test siRNA data was normalized relative to the mean of negative control samples (set as 0% effect) and the mean of positive control samples (set as 100% effect). Three standard deviations of the negative control samples were used as a cutoff to define screen actives. As for manual curation of the literature to identify genes without previous functional validation associated with cancer, we based results on an extensive search of PubMed. Each gene with studies where it was deliberately overexpressed, suppressed, or mutated and resulted an *in vitro* change in the phenotype of cancer cell lines was annotated as functionally validated. Full annotated lists are provided in the supplement.

## References

1. Pon JR, Marra MA: Driver and passenger mutations in cancer. Annu Rev Pathol 2015, 10:25–50.

2. Loganathan SK, Schleicher K, Malik A, Quevedo R, Langille E, Teng K, Oh RH, Rathod B, Tsai R, Samavarchi-Tehrani P, et al: **Rare driver mutations in head and neck squamous cell carcinomas converge on NOTCH signaling**. Science 2020, 367:1264–1269.

3. Scholl C, Frohling S: Exploiting rare driver mutations for precision cancer medicine. Curr Opin Genet Dev 2019, 54:1–6.

4. Armenia J, Wankowicz SAM, Liu D, Gao J, Kundra R, Reznik E, Chatila WK, Chakravarty D, Han GC, Coleman I, et al: **The long tail of oncogenic drivers in prostate cancer**. Nat Genet 2018, 50:645–651.

5. Elman JS, Ni TK, Mengwasser KE, Jin D, Wronski A, Elledge SJ, Kuperwasser C: Identification of FUBP1 as a Long Tail Cancer Driver and Widespread Regulator of Tumor Suppressor and Oncogene Alternative Splicing. Cell Rep 2019, 28:3435–3449 e3435.

6. Consortium I-TP-CAoWG: **Pan-cancer analysis of whole genomes**. Nature 2020, 578:82–93.

7. Bailey MH, Tokheim C, Porta-Pardo E, Sengupta S, Bertrand D, Weerasinghe A, Colaprico A, Wendl MC, Kim J, Reardon B, et al: **Comprehensive Characterization of Cancer Driver Genes and Mutations**. Cell 2018, 173:371–385 e318.

8. Nitsch D, Gonçalves JP, Ojeda F, de Moor B, Moreau Y: Candidate gene prioritization by network analysis of differential expression using machine learning approaches. BMC Bioinformatics 2010, 11:460.

9. Lee I, Blom UM, Wang PI, Shim JE, Marcotte EM: Prioritizing candidate disease genes by network-based boosting of genome-wide association data. Genome Res 2011, 21:1109–1121.

10. Erten S, Bebek G, Ewing RM, Koyuturk M: DADA: Degree-Aware Algorithms for Network-Based Disease Gene Prioritization. BioData Min 2011, 4:19.

11. Erten S, Bebek G, Koyuturk M: Vavien: an algorithm for prioritizing candidate disease genes based on topological similarity of proteins in interaction networks. J Comput Biol 2011, 18:1561–1574.

12. Cao M, Zhang H, Park J, Daniels NM, Crovella ME, Cowen LJ, Hescott B: Going the distance for protein function prediction: a new distance metric for protein interaction networks. PLoS One 2013, 8:e76339.

13. Cao M, Pietras CM, Feng X, Doroschak KJ, Schaffner T, Park J, Zhang H, Cowen LJ, Hescott BJ: **New directions for diffusion-based network prediction of protein function: incorporating pathways with confidence**. Bioinformatics 2014, 30:i219–227.

14. Cowen L, Ideker T, Raphael BJ, Sharan R: **Network propagation: a universal amplifier of genetic associations**. Nat Rev Genet 2017, 18:551–562.

15. Köhler S, Bauer S, Horn D, Robinson PN: Walking the interactome for prioritization of candidate disease genes. Am J Hum Genet 2008, 82:949–958.

16. Vanunu O, Magger O, Ruppin E, Shlomi T, Sharan R: **Associating genes and protein complexes with disease via network propagation**. PLoS Comput Biol 2010, 6:e1000641.

17. Paull EO, Carlin DE, Niepel M, Sorger PK, Haussler D, Stuart JM: Discovering causal pathways linking genomic events to transcriptional states using Tied Diffusion Through Interacting Events (TieDIE). Bioinformatics 2013, 29:2757–2764.

18. Singh-Blom UM, Natarajan N, Tewari A, Woods JO, Dhillon IS, Marcotte EM: Prediction and validation of gene-disease associations using methods inspired by social network analyses. PLoS One 2013, 8:e58977.

19. Ruffalo M, Koyuturk M, Sharan R: Network-Based Integration of Disparate Omic Data To Identify “Silent Players” in Cancer. PLoS Comput Biol 2015, 11:e1004595.

20. Shnaps O, Perry E, Silverbush D, Sharan R: **Inference of Personalized Drug Targets Via Network Propagation**. Pac Symp Biocomput 2016, 21:156–167.

21. Hofree M, Shen JP, Carter H, Gross A, Ideker T: **Network-based stratification of tumor mutations**. Nat Methods 2013, 10:1108–1115.

22. Reyna MA, Leiserson MDM, Raphael BJ: **Hierarchical HotNet: identifying hierarchies of altered subnetworks**. Bioinformatics 2018, 34:i972–i980.

23. Vandin F, Upfal E, Raphael BJ: Algorithms for detecting significantly mutated pathways in cancer. J Comput Biol 2011, 18:507–522.

24. Leiserson MD, Vandin F, Wu HT, Dobson JR, Eldridge JV, Thomas JL, Papoutsaki A, Kim Y, Niu B, McLellan M, et al: **Pan-cancer network analysis identifies combinations of rare somatic mutations across pathways and protein complexes**. Nat Genet 2015, 47:106–114.

25. McFarland CD, Mirny LA, Korolev KS: Tug-of-war between driver and passenger mutations in cancer and other adaptive processes. Proc Natl Acad Sci U S A 2014, 111:15138–15143.

26. Castro-Giner F, Ratcliffe P, Tomlinson I: **The mini-driver model of polygenic cancer evolution**. Nat Rev Cancer 2015, 15:680–685.

27. Nussinov R, Tsai CJ: ’Latent drivers’ expand the cancer mutational landscape. Curr Opin Struct Biol 2015, 32:25–32.

28. McFarland CD, Yaglom JA, Wojtkowiak JW, Scott JG, Morse DL, Sherman MY, Mirny LA: **The Damaging Effect of Passenger Mutations on Cancer Progression**. Cancer Res 2017, 77:4763–4772.

29. Kumar S, Warrell J, Li S, McGillivray PD, Meyerson W, Salichos L, Harmanci A, Martinez-Fundichely A, Chan CWY, Nielsen MM, et al: **Passenger Mutations in More Than 2,500 Cancer Genomes: Overall Molecular Functional Impact and Consequences.** Cell 2020, 180:915–927 e916.

30. Szklarczyk D, Gable AL, Lyon D, Junge A, Wyder S, Huerta-Cepas J, Simonovic M, Doncheva NT, Morris JH, Bork P, et al: STRING v11: protein-protein association networks with increased coverage, supporting functional discovery in genome-wide experimental datasets. Nucleic Acids Res 2019, 47:D607–D613.

31. Hwang S, Kim CY, Yang S, Kim E, Hart T, Marcotte EM, Lee I: **HumanNet v2: human gene networks for disease research**. Nucleic Acids Res 2019, 47:D573–D580.

32. Tsherniak A, Vazquez F, Montgomery PG, Weir BA, Kryukov G, Cowley GS, Gill S, Harrington WF, Pantel S, Krill-Burger JM, et al: **Defining a Cancer Dependency Map**. Cell 2017, 170:564–576 e516.

33. Krzywinski M, Schein J, Birol I, Connors J, Gascoyne R, Horsman D, Jones SJ, Marra MA: **Circos: an information aesthetic for comparative genomics**. Genome Res 2009, 19:1639–1645.

34. Sanchez-Vega F, Mina M, Armenia J, Chatila WK, Luna A, La KC, Dimitriadoy S, Liu DL, Kantheti HS, Saghafinia S, et al: **Oncogenic Signaling Pathways in The Cancer Genome Atlas**. Cell 2018, 173:321–337 e310.

35. Reimand J, Isserlin R, Voisin V, Kucera M, Tannus-Lopes C, Rostamianfar A, Wadi L, Meyer M, Wong J, Xu C, et al: **Pathway enrichment analysis and visualization of omics data using g:Profiler, GSEA, Cytoscape and EnrichmentMap**. Nat Protoc 2019, 14:482–517.

36. Herrero-Gonzalez S, Di Cristofano A: New routes to old places: PIK3R1 and PIK3R2 join PIK3CA and PTEN as endometrial cancer genes. Cancer Discov 2011, 1:106–107.

37. Pan F, Zhang J, Tang B, Jing L, Qiu B, Zha Z: The novel circ_0028171/miR-218-5p/IKBKB axis promotes osteosarcoma cancer progression. Cancer Cell Int 2020, 20:484.

38. Torres HA, Shigle TL, Hammoudi N, Link JT, Samaniego F, Kaseb A, Mallet V: **The oncologic burden of hepatitis C virus infection: A clinical perspective**. CA Cancer J Clin 2017, 67:411–431.

39. Haggstrom C, Van Hemelrijck M, Zethelius B, Robinson D, Grundmark B, Holmberg L, Gudbjornsdottir S, Garmo H, Stattin P: **Prospective study of Type 2 diabetes mellitus, anti-diabetic drugs and risk of prostate cancer**. Int J Cancer 2017, 140:611–617.

40. Shlomai G, Neel B, LeRoith D, Gallagher EJ: **Type 2 Diabetes Mellitus and Cancer: The Role of Pharmacotherapy**. J Clin Oncol 2016, 34:4261–4269.

41. Tagaya Y, Gallo RC: **The Exceptional Oncogenicity of HTLV-1**. Front Microbiol 2017, 8:1425.

42. McFarland JM, Ho ZV, Kugener G, Dempster JM, Montgomery PG, Bryan JG, Krill-Burger JM, Green TM, Vazquez F, Boehm JS, et al: **Improved estimation of cancer dependencies from large-scale RNAi screens using model-based normalization and data integration**. Nat Commun 2018, 9:4610.

43. Yang L, Chen E, Goodison S, Sun Y: An efficient and effective method to identify significantly perturbed subnetworks in cancer. Nat Comput Sci 2021, 1:79–88.

44. Hristov BH, Singh M: Network-Based Coverage of Mutational Profiles Reveals Cancer Genes. Cell Syst 2017, 5:221–229 e224.

45. Lever J, Zhao EY, Grewal J, Jones MR, Jones SJM: CancerMine: a literature-mined resource for drivers, oncogenes and tumor suppressors in cancer. Nat Methods 2019, 16:505–507.

46. Ellrott K, Bailey MH, Saksena G, Covington KR, Kandoth C, Stewart C, Hess J, Ma S, Chiotti KE, McLellan M, et al: **Scalable Open Science Approach for Mutation Calling of Tumor Exomes Using Multiple Genomic Pipelines**. Cell Syst 2018, 6:271–281 e277.

47. Wang K, Li M, Hakonarson H: ANNOVAR: functional annotation of genetic variants from high-throughput sequencing data. Nucleic Acids Res 2010, 38:e164.

48. Durinck S, Spellman PT, Birney E, Huber W: Mapping identifiers for the integration of genomic datasets with the R/Bioconductor package biomaRt. Nat Protoc 2009, 4:1184–1191.

49. Wang Q, Armenia J, Zhang C, Penson AV, Reznik E, Zhang L, Minet T, Ochoa A, Gross BE, Iacobuzio-Donahue CA, et al: **Unifying cancer and normal RNA sequencing data from different sources**. Sci Data 2018, 5:180061.

50. Ramakrishnan SR, Vogel C, Kwon T, Penalva LO, Marcotte EM, Miranker DP: Mining gene functional networks to improve mass-spectrometry-based protein identification. Bioinformatics 2009, 25:2955–2961.

51. Zhou DY, Bousquet O, Lal TN, Weston J, Scholkopf B: **Learning with local and global consistency**. Advances in Neural Information Processing Systems 16 2004, 16:321–328.

52. Langville ANaM, Carl D.: **Deeper Inside PageRank**. Internet Mathematics 2003, 1.

53. Tate JG, Bamford S, Jubb HC, Sondka Z, Beare DM, Bindal N, Boutselakis H, Cole CG, Creatore C, Dawson E, et al: **COSMIC: the Catalogue Of Somatic Mutations In Cancer**. Nucleic Acids Res 2019, 47:D941–D947.

54. Lawrence MS, Stojanov P, Polak P, Kryukov GV, Cibulskis K, Sivachenko A, Carter SL, Stewart C, Mermel CH, Roberts SA, et al: **Mutational heterogeneity in cancer and the search for new cancer-associated genes**. Nature 2013, 499:214–218.

55. Jiao X, Sherman BT, Huang da W, Stephens R, Baseler MW, Lane HC, Lempicki RA: **DAVID-WS: a stateful web service to facilitate gene/protein list analysis**. Bioinformatics 2012, 28:1805–1806.

56. Shannon P, Markiel A, Ozier O, Baliga NS, Wang JT, Ramage D, Amin N, Schwikowski B, Ideker T: **Cytoscape: a software environment for integrated models of biomolecular interaction networks**. Genome Res 2003, 13:2498–2504.

